# Preliminary characterization of CbcS from *Geobacter sulfurreducens*’ Cbc4 complex: a putative novel respiratory pathway

**DOI:** 10.1101/2025.04.14.648717

**Authors:** Mafalda V. Fernandes, Jorge M. A. Antunes, Carlos A. Salgueiro, Leonor Morgado

## Abstract

Electroactive bacteria mediate electron exchange with external surfaces through a process known as extracellular electron transfer (EET). A key step in EET is the transfer of electrons from the menaquinone pool to inner membrane-associated quinone-cytochrome *c* oxidoreductase complexes, which subsequently relay electrons to periplasmic redox partners. Gene knockout and proteomic analyses have identified several critical components involved in EET in *Geobacter sulfurreducens*, including six inner membrane oxidoreductase gene clusters. Of these, three - CbcL, ImcH, and CbcBA - have been linked to specific respiratory pathways depending on the redox potential of the terminal electron acceptor.

Cbc4 is one of the other inner membrane oxidoreductase complexes and is composed by a membrane-anchored tetraheme *c*-type cytochrome (CbcS), an iron–sulfur protein containing four [4Fe-4S] clusters (CbcT), and an integral membrane subunit (CbcU). In this study, the sequence and AlphaFold model of CbcS were analyzed and its cytochrome domain was produced, and structurally and functionally characterized. Nuclear Magnetic Resonance spectroscopy data validated the hemecore arrangement predicted by AlphaFold, showing that, despite the differences in axial ligands (CbcS has four bis-histidine low-spin hemes), CbcS hemecore is homologous to CymA and NrfH from *Shewanella* and *Desulfovibrio* species, respectively. Potentiometric titrations showed that CbcS redox active window superimposes with the ones of its putative redox partners from the triheme periplasmic cytochrome family PpcA-E; however, electron transfer reactions monitored by NMR revealed that CbcS is able to transfer electrons to PpcA. Furthermore, NMR redox titrations allowed to identify heme IV as the exit gate for electrons. Together, these findings contribute to the understanding of the molecular mechanisms of EET and provide insights on a putative new respiratory pathway in *G. sulfurreducens*.

## Introduction

Microorganisms, particularly anaerobic bacteria, have played a fundamental role in Earth’s biogeochemical cycles for nearly four billion years, shaping the distribution and availability of essential elements ^1^. Early anaerobic bacteria thrived in oxygen-free environments by utilizing minerals as electron donors and transferring electrons to acceptors such as nitrate or sulfate. With the emergence of an oxidizing atmosphere, some bacterial species evolved to incorporate oxygen into their metabolic pathways ^1,2^. Among these microorganisms, there is a type of bacteria that are especially important since they contribute to the cycling of metals by coupling the oxidation of organic matter to the reduction of metals. In conventional respiratory pathways, the reduction occurs inside the cell, but these bacteria, dissimilatory metal-reducing bacteria (DMRB), have developed electron transfer chains that are capable of transferring electrons across the membranes to reduce metals externally ^3,4^. This unique capability has positioned DMRB as key players in bioremediation, enabling the reduction of toxic and radioactive metals. DMRB are also classified as exoelectrogens - microorganisms that transfer electrons to external surfaces, including electrodes, thereby facilitating the development of bioelectrochemical systems ^5^. One notable application is microbial fuel cells (MFCs), which harness microbial metabolism to convert organic matter into electricity. MFCs hold promise for sustainable energy production while simultaneously contributing to wastewater treatment, aligning with global sustainability efforts ^6,7^.

As environmental challenges such as pollution, climate change, and access to clean water intensify, understanding and optimizing electron transfer pathways in DMRB becomes increasingly crucial. The discovery of electricity-generating bacteria dates back over a century ^8^. Since then, *Geobacter* species have demonstrated their effectiveness in bioremediation, particularly in groundwater decontamination ^5^, and their capacity for extracellular electron transfer (EET) has positioned them as prime candidates for bioelectrochemical technologies ^1^. Similarly, *Shewanella* species can conserve energy through Fe(III) and Mn(IV) reduction, making *Geobacter* and *Shewanella* the most extensively studied electroactive microorganisms and model systems for investigating EET mechanisms ^9,10^.

During EET, electrons generated from the oxidation of nutrients in the cytoplasm are relayed through a series of cytochromes located in the inner membrane, periplasm, and outer membrane, ultimately reaching terminal electron acceptors outside the cell ^9^. The current model of EET pathways in *Geobacter* is supported by integrated proteomic and genomic analysis ^2,11^, and suggests that electrons derived from nicotinamide adenine dinucleotide (NADH) are first transferred to the menaquinone pool in the inner membrane via NADH dehydrogenase. From the menaquinone pool, electrons flow through quinone-cytochrome *c* oxidoreductases located in the inner membrane and are subsequently transferred to the triheme cytochromes family (PpcA, PpcB, PpcC, PpcD, and PpcE) in the periplasm. From there, the electrons are relayed to proteins anchored to the outer membrane, ultimately being transferred to extracellular electron acceptors such as Fe(III) oxides, Mn(IV) oxides, or electrodes _12–16_.

The genome of *Geobacter sulfurreducens* encodes six quinone-cytochrome *c* oxidoreductases located in the inner membrane. These include four cytochrome *bc* (Cbc) complexes (Cbc3 to Cbc6), CbcL with both *c*-type and *b*-type hemes and ImcH with only *c*-type hemes ^13–15^. The Cbc complexes are predicted to facilitate electron transfer between the menaquinone pool and the periplasmic cytochromes, while also generating a proton gradient across the inner membrane that drives ATP synthesis. These complexes are highly conserved in *Geobacter* species due to their essential role ^11,17^. Gene-knockout and proteomic studies have already established a role for CbcL, ImcH, and CbcBA from Cbc5, and showed that the deletion of these proteins severely impairs the ability of the bacteria to reduce certain electron acceptors ^13–15^. However, no specific role for the remaining complexes has been attributed so far.

One of these complexes is Cbc4, that is formed by an anchored tetraheme *c*-type cytochrome (CbcS - GSU0068), an iron-sulfur protein containing four [4Fe-4S] centers (CbcT - GSU0069), and an integral membrane subunit (CbcU - GSU0070). The *cbcS* gene is upregulated in the presence of Fe(III) oxides but downregulated in the presence of Mn(IV) oxides. Notably, mutant strains lacking the *cbcS* gene exhibit growth similar to that of wild-type strains, indicating that CbcS is not essential for Fe(III) and Mn(IV) reduction. Both *cbcU* and *cbcT* are upregulated in the presence of insoluble metal oxides ^12^.

This study focuses on a primary characterization of the structure, function, and role of cytochrome CbcS within the electron transfer network of *G. sulfurreducens*, providing insights into the intricate molecular strategies employed by these versatile bacteria.

### Experimental procedures Bioinformatics analysis

The DNA sequence of the gene coding for CbcS (*cbcS*, GSU0068, UNIPROT Q74H25) was extracted from *G. sulfurreducens* PCA genome from the Kyoto Encyclopedia of Genes and Genomes database ^18–20^. The software SignalP 6.0 ^21^ and DeepTMHMM 1.0 ^22^ were used for prediction of the presence of a signal peptide and of transmembrane helices, respectively. A structural model of CbcS was generated with AlphaFold 3 ^23^ using as input the amino acid sequence and four *c*-type hemes. The model was analyzed with PyMOL (The PyMOL Molecular Graphics System, Version 2.5.7 Schrödinger, LLC) and the images were prepared with ChimeraX 1.9 ^24–26^.

### Cloning

CbcS DNA sequence was amplified from *G. sulfurreducens* genomic DNA using Phusion High-Fidelity DNA Polymerase (Thermo Fisher Scientific) and the primers listed in Table S1. Two constructs were prepared – one including residues 30-160 (ST-CbcS_pp_) and the other residues 59-160 (ST-CbcS_cyt_) – both with an N-terminal Strep-tag to facilitate the purification process. PCR products were purified with the NZYGelpure kit (NZYTech) and, using the restriction-free cloning method ^27,28^, the genes were then inserted into vector pVA203 ^29,30^. DpnI (NZYTech) was used for template DNA digestion before transformation in competent *Escherichia coli* DH5α cells. Positive clones were selected by colony screening via PCR using NZYTaq II DNA Polymerase (NZYTech) monitored by agarose gel electrophoresis and cultured in liquid medium for plasmid propagation. Plasmid DNA was purified using the NZYMiniprep kit (NZYTech), and their concentration and purity were assessed with a NanoDrop spectrophotometer ND-100 (Thermo Fisher Scientific). DNA sequencing was performed by STABVida (Caparica, Portugal).

A Tobacco Etch Virus (TEV) protease ^31^ cleavage site was further introduced in plasmid ST-CbcS_cyt_ to allow removal of the Strep-tag, using a strategy based on the Q5 Site-Directed Mutagenesis kit (New England Biolabs) and the primers listed in Table S1. The PCR product was treated with DpnI, purified with the NZYGelpure kit and the DNA concentration was measured spectrophotometrically. The following step involved incubation with T4 Polynucleotide Kinase (NZYTech) and T4 DNA Ligase (NZYTech) at 30°C for 1 hour to allow circularization. The DNA was then transformed into *E. coli* DH5α cells and positive clones selected and sequenced as described above.

### Protein expression and purification

*E. coli* BL21 (DE3) cells were co-transformed with two plasmids: pEC86, which encodes the cytochrome *c* maturation gene cluster and confers chloramphenicol resistance, and pVA203-CbcS, which harbors the *lac* promoter, an ampicillin resistance gene, the OmpA signal peptide for periplasmic translocation and the gene coding for CbcS ^30,32,33^. Co-expression of pEC86 facilitates proper heme cofactor incorporation into the target protein.

Transformed *E. coli* colonies carrying the CbcS construct were inoculated into 50 mL of 2×YT medium supplemented with 100 µg.mL^-1^ of ampicillin and 34 µg.mL^-1^ of chloramphenicol and incubated overnight at 30°C at 200 rpm. The following day, 20 mL of the overnight culture were used to inoculate 1 L of 2×YT medium which were then incubated at 30°C at 180 rpm. When the culture reached an optical density at 600 nm of ∼1.5, protein expression was induced with 10 μM of isopropyl β-D-1-thiogalactopyranoside (IPTG) and incubated for 2 hours. Cells were harvested by centrifugation at 6,400 *xg* for 10 minutes at 6°C, and the resulting pellets were resuspended in buffer W (100 mM Tris-HCl pH 8, 150 mM NaCl, 1 mM EDTA).

Cell lysis was performed through three freeze-thaw cycles, in the presence of 1 mM phenylmethylsulphonyl fluoride (PMSF), 2 mM benzamidine, and DNase I (Roche). Further disruption was achieved using 20 cycles of ultrasonication (2 minutes on, 1 minute off) at 65% amplitude using an ultrasonic homogenizer (Branson).. Insoluble debris was removed by centrifugation at 49,000 *xg* for 90 minutes at 6°C. The clarified lysate, containing soluble CbcS, was subjected to affinity chromatography using a 5 mL Strep-Tactin® XT 4Flow® column (IBA) pre-equilibrated with buffer W. After washing with 30 mL of buffer W, bound protein was eluted with ∼30 mL of buffer BXT (buffer W with 50 mM biotin).

Protein-containing fractions were analyzed by SDS-PAGE using a Tris-tricine buffer system ^34^ and stained with BlueSafe (NZYTech). CbcS fractions were concentrated using an Amicon Ultra-4 ultrafiltration device (3 kDa molecular weight cutoff, Merck) and further purified by size-exclusion chromatography on a Superdex™ 75 Increase 10/300 GL column (Cytiva) equilibrated with 100 mM sodium phosphate buffer pH 8. In the case of the construct used for tag removal, after the first purification step, the fractions containing ST-CbcS_cyt_ were dialyzed against 50 mM Tris HCl buffer pH 8 with 150 mM NaCl, 0.5 mM EDTA and 1 mM DTT in the presence of TEV protease ^31^ (ratio TEV:CbcS 1:5 mg) for overnight cleavage at 4 °C. After cleavage, the protein was applied to the Strep-Tactin® XT 4Flow® column to separate any uncleaved protein and further concentrated by ultrafiltration before being applied to the size-exclusion column (that efficiently separated CbcS_cyt_ from TEV). Chromatographic purification steps were performed using ÄKTA Pure or ÄKTA Prime Plus system. The yields for St-CbcS_pp_, ST-CbcS_cyt_ and CbcS_cyt_ were 0.19, 0.52 and 0.21 mg per liter of culture, respectively.

Final protein purity was assessed by SDS-PAGE, and the molecular weight of CbcS_cyt_ was confirmed by matrix-assisted laser desorption/ionization time-of-flight mass spectrometry (MALDI-TOF/TOF MS) at the Mass Spectrometry Unit (UniMS), ITQB/iBET, Oeiras, Portugal.

PpcA from *G. sulfurreducens* was produced as previously described ^29^.

### Protein and heme quantification and molar extinction coefficient determination

UV-Visible absorption spectra of purified CbcS were recorded in the oxidized as-purified and reduced states on a Thermo Scientific Evolution 201 spectrophotometer at room temperature. Spectra were acquired over a wavelength range of 750–250 nm in 1 cm path length quartz cuvettes (Hellma). The protein was reduced by adding sodium dithionite crystals. Protein concentration was determined using the Pierce™ Modified Lowry Protein Assay Kit (Thermo Fisher Scientific), using horse heart cytochrome *c* as the standard protein for the calibration curve, and it was used to determine CbcS molar extinction coefficient. All the subsequent protein concentrations were determined using the calculated molar extinction coefficient at the absorption value of 552 nm in the reduced state (α-band). The heme content was determined by the pyridine hemochromogen assay by measuring the absorbance at 550 nm (ε_550nm_ = 30.27 mM^-1^.cm^-1^ for *c*-type cytochromes) of 1.6 µM CbcS solution prepared in 75 mM of NaOH/25% pyridine and reduced by the addition of sodium dithionite crystals ^35^.

### Potentiometric titrations followed by UV-visible spectroscopy

Redox titrations of ST-CbcS_cyt_ were conducted under anaerobic conditions within an anaerobic glovebox (MBraun) under an argon atmosphere. The experiments were performed using ∼8 µM CbcS in 80 mM sodium phosphate buffer with NaCl (250 mM ionic strength) at pH 7 and pH 8. The solution potential was monitored using a platinum ring electrode coupled with an AgCl/Ag reference electrode (Mettler Toledo), and values were converted to the standard hydrogen electrode (SHE) reference. Calibration of the electrode was performed at the beginning and end of each titration using freshly prepared saturated quinhydrone solutions at pH 7 and pH 4. The titrations were conducted in a cuvette holder (OceanOptics) inside the anaerobic glovebox, which was connected to a UV-Visible spectrophotometer (Thermo Scientific Evolution 300) via optical fibers (Horiba). The solution temperature was maintained at 15°C using a circulating bath (VWR). A mixture of the following redox mediators was added to a final concentration of 1 µM, to facilitate electron exchange between the redox centers and the electrode: 1,2-napthoquinone (E^0’^ = +143 mV), trimethylhydroquinone (E^0’^ = +115 mV), phenazine methosulfate (E^0’^ = +80 mV), phenazine ethosulfate (E^0’^ = +55 mV), gallocyanin (E^0’^ = +21 mV), methylene blue (E^0’^ = +11 mV), indigo tetrasulfonate (E^0’^ = -30 mV), indigo trisulfonate (E^0’^ = -70 mV), indigo disulfonate (E^0’^ = -120 mV), 2-hydroxy-1,4-naphtoquinone (E^0’^ = -145 mV), antraquinone-2,6-disulfonate (E^0’^ = -185 mV), antraquinone-2-sulfonate (E^0’^ = -225 mV), safranine (E^0’^ = -280 mV), neutral red (E^0’^ = -325 mV), benzyl viologen(E^0’^ = -345 mV), diquat (E^0’^ = -350 mV) and methyl viologen (E^0’^ = -440 mV) ^36,37^. Sample reduction was achieved using sodium dithionite, while potassium ferricyanide was used as the oxidizing agent.

The reduced fraction was quantified by integrating the absorbance area above the isosbestic points at 549 and 568 nm, following the method described by ^38^. The apparent macroscopic midpoint reduction potential (*E*_*app*_) was determined by fitting the experimental data to a nonlinear model considering four sequential one-electron Nernst equations. Data analysis and fitting were performed using OriginPro 8.

### NMR spectroscopy

Nuclear Magnetic Resonance (NMR) experiments were performed using a Bruker Avance III 600 MHz spectrometer equipped with a triple resonance cryoprobe (TCI). The proton chemical shifts were calibrated using the water signal as internal reference ^39^. Spectral data were processed using TopSpin 3.6.5™ (Bruker BioSpin, Karlsruhe, Germany) and analyzed with Sparky 3.115.

#### Sample Preparation

Samples were subjected to three lyophilization cycles in 10 mM sodium phosphate pH 8 buffer and subsequently resuspended in either ^2^H2O or 80 mM sodium phosphate pH 8 buffer (250 mM ionic strength) prepared in ^2^H2O. Due to precipitation at lower buffer concentrations, ST-CbcS_pp_ samples were prepared in 50 mM sodium phosphate pH 8 buffer. Reported pH values were not corrected for isotope effects.

For the preparation of reduced CbcS, the NMR tube was sealed with a gas-tight rubber cap, and the sample was purged with gaseous hydrogen. A catalytic amount of hydrogenase from *Desulfovibrio vulgaris* was added to facilitate reduction. For experiments in intermediate redox states and electron transfer experiments, the hydrogen was removed by argon flushing after reduction. For 2D ^1^H-EXchange SpectroscopY (2D EXSY), controlled amounts of air were added to the NMR tube with a Hamilton gas-tight syringe.

#### Initial NMR experiments in the oxidized state

One-dimensional ^1^H NMR spectra of oxidized CbcS variants were acquired in ^2^H2O to evaluate structural changes. ST-CbcS_cyt_ and CbcS_cyt_ samples were prepared in 80 mM sodium phosphate pH 8 buffer (250 mM ionic strength), while ST-CbcS_pp_ was prepared in 50 mM sodium phosphate pH 8 buffer. Spectra were recorded at 25°C using the *zgpr* pulse sequence with a sweep width of 80 ppm and 4096 scans for ST-CbcS_pp_, 256 scans for ST-CbcS_cyt_, and 128 scans for CbcS_cyt_.

#### NMR experiments in the fully reduced state

For heme resonance assignments in the fully reduced state, a sample of CbcS_cyt_ was prepared in 80 mM sodium phosphate buffer pH 8.1 (250 mM ionic strength). 2D ^1^H TOCSY (*dipsy2phpr*) and 2D ^1^H NOESY (*noesyphpr*) experiments were acquired with mixing times of 60 ms and 80 ms, respectively, a sweep width of 20 ppm using 2k (t_2_) x 512 (t_1_) data points and 128 scans. Spectra were acquired at 9°C, 15°C or 25°C.

#### NMR experiments in intermediate redox states

2D EXSY spectra were acquired in a 1mM CbcS_cyt_ sample in 80 mM sodium phosphate pH 8.1 buffer (250 mM ionic strength) at 9°C and different oxidation states. Initial tests were performed at lower concentration ∼ 70µM and different temperatures (9°C, 15°C and 25°C) ^40,41^. The 2D ^1^H EXSY (*noesyphpr*) spectra were acquired with a mixing time of 25 ms, a sweep width of 76 ppm using 2k (t_2_) x 256 (t_1_) data points, and 256 scans. 1D ^1^H NMR spectra were obtained before and after each 2D EXSY spectrum to monitor any changes in the oxidation state of the sample during the 2D experiment.

#### Electron transfer experiments

The electron transfer reaction between reduced CbcS and oxidized PpcA was monitored by ^1^H NMR as previously described for other cytochromes ^42^. To follow the electron transfer reaction of CbcS and PpcA, a sample of 45 µM ST-CbcS_cyt_ (550 µL) and a sample of PpcA at 90 µM (550 µL) both in 10 mM sodium phosphate buffer (pH 8.1) were lyophilized. CbcS sample was resuspended in 550 µL of ^2^H_2_O and transferred to a 5 mm diameter NMR tube and PpcA was resuspended in 30 µL of ^2^H_2_O. CbcS reduction occurred as described before and increasing equivalents of PpcA were sequentially added under anaerobic conditions inside a glovebox. The 1D ^1^H NMR spectra were acquired with 512 scans and a sweep width of 80 ppm.

## Results and discussion

### CbcS primary sequence and structural model analysis

The analysis of CbcS primary sequence revealed that the first 29 residues form a membrane anchor and that four *c*-type heme binding motifs (CXXCH) are present (Figure S1). Different hemecore arrangements have been previously described for tetraheme cytochromes. In the Small Tetraheme Cytochrome (STC) from *Shewanella oneidensis* the hemes form a chain-like configuration, while in cytochrome *c*_3_ from *Desulfovibrio vulgaris* the hemes are arranged in a circular structure (Figure 1 and Figure S2) ^43,44^. Both types of cytochromes have all four hemes coordinated by two histidine residues, and the relative positions of the heme binding motifs and the respective distal axial ligands in their primary sequence lead to important modifications both at the level of the heme core architecture and at the level of the general folding ^41^.

**Figure 1.**
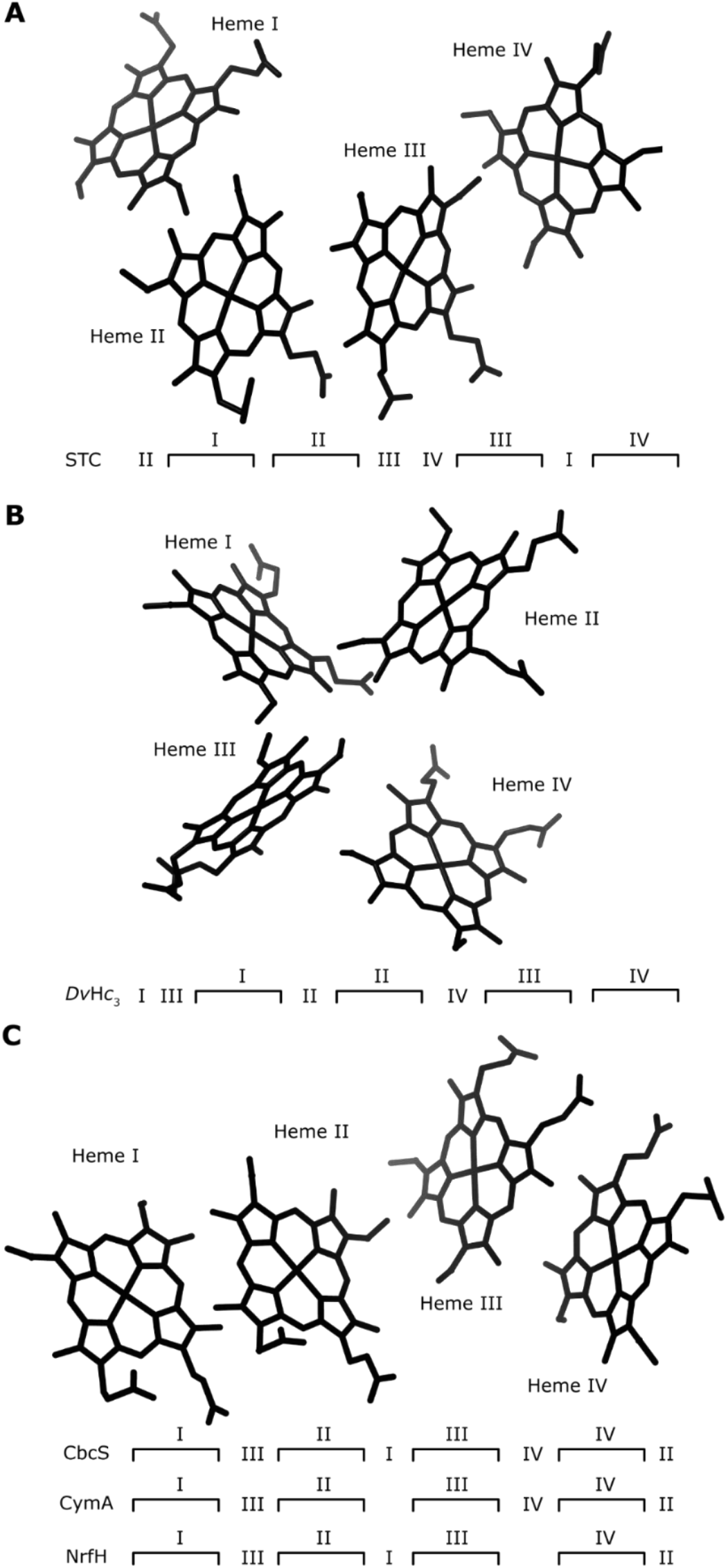
Hemecore arrangement and order of heme binding sites and distal ligand for different tetraheme cytochromes. **(A)** Small tetraheme cytochrome *c* (STC) from *Shewanella frigidimarina* (PBD 2K3V) ^43^. **(B)** Cytochrome *c*_3_ from *Desulfovibrio vulgaris* Hildenborough (PDB 1A2I) ^44^. **(C)** CbcS (AlphaFold3 prediction). CymA from *S. frigidimarina* and NrfH from *D. vulgaris* show a similar hemecore arrangement but differences in the distal ligands. In NrfH, the heme IV distal ligand is a lysine from NrfA from the same complex ^45^. Previous studies on CymA and its structural model prediction (Figure S1) show the absence of a distal ligand for heme I ^46^.

NrfH, a membrane anchored small protein of the cytochrome *c* nitrite reductase NrfHA complex from *D. vulgaris*, is also a tetraheme cytochrome that belongs to the NapC/NirT family ^45^. In the crystal structure of the NrfHA complex, NrfH hemes II and III are bis-His coordinated, heme I is proximally coordinated by a methionine and distally by an aspartate (thus having high-spin characteristics), and heme IV is coordinated by a histidine and has a lysine from NrfA as a distal ligand (Figure S2). Similarly to STC, NrfH hemes are arranged in an overall chain configuration. According to sequence alignments, CymA, a crucial quinol oxidoreductase in *Shewanella*’s EET mechanisms, is also a member of the NapC/NirT family ^10^. CymA’s tridimensional structure was never determined, but NMR studies showed that it binds three low-spin hemes with bis-histidine coordination, and one high-spin spin heme with histidine-water coordination ^46,47^. AlphaFold predictions also corroborate this hypothesis (Figure S2). Attempts to produce a soluble form of CymA, truncated at residue 35 for removal of the membrane anchor, led to the production of a tetraheme cytochrome with four low-spin hemes ^48^. This phenomenon is likely attributed to the interaction of histidine 38 with heme I. In the native form, histidine 38 is positioned away from heme I, but truncation of the transmembrane domain may increase its flexibility, allowing it to coordinate the heme at its available axial binding position. An analysis of the possible distal ligands for CbcS hemes, together with its AlphaFold structural model (Figure S1) predicts that all four hemes are bis-histidine coordinated, with a hemecore arrangement similar to the one of NrfH and CymA (Figure 1). In contrast with what it’s observed for CymA and NrfH, this suggests that CbcS heme groups are low-spin.

The AlphaFold model of CbcS also predicts the existence of an unstructured linker domain (residues 30–58) between the transmembrane anchor and the cytochrome domain (Figure S1). This linker might have a role in providing flexibility to CbcS when interacting with other Cbc4 proteins or its upstream redox partners. Based on these observations, different constructs were strategically designed to produce CbcS in a soluble form, while minimizing structural alterations resulting from residue deletions.

### CbcS cytochrome domain maintains its global fold in different constructs

CbcS was produced with an N-terminal cleavable Strep-tag in two constructs: ST-CbcS_pp_ that expresses the cytochrome domain together with the linker domain, and ST-CbcS_cyt_ that expresses only the cytochrome domain. Both proteins were produced, and pure samples were obtained after one step of affinity chromatography followed by a step of size exclusion chromatography (Figure 2A) with yields of 0.19 and 0.52 mg per liter of culture for ST-CbcS_pp_ and ST-CbcS_cyt_, respectively. However, ST-CbcS_pp_ revealed to be less stable since some precipitation occurred during buffer exchange, as well as through time. As heme resonances are extremely sensitive to structural changes, particularly in the oxidized state due to paramagnetic effects, both proteins were analyzed by 1D ^1^H NMR in the oxidized state (Figure S3). Resonances match for both constructs (slight differences are due to differences in buffer conditions), which indicates that there are no structural changes in the absence of the linker domain. Since the spectra were identical, but a lower protein yield was obtained for the larger construct, all the subsequent experiments were performed with ST-CbcS_cyt_. Also, while the linker domain may be essential within the Cbc4 complex, the functional domain of CbcS resides in the cytochrome region.

**Figure 2.**
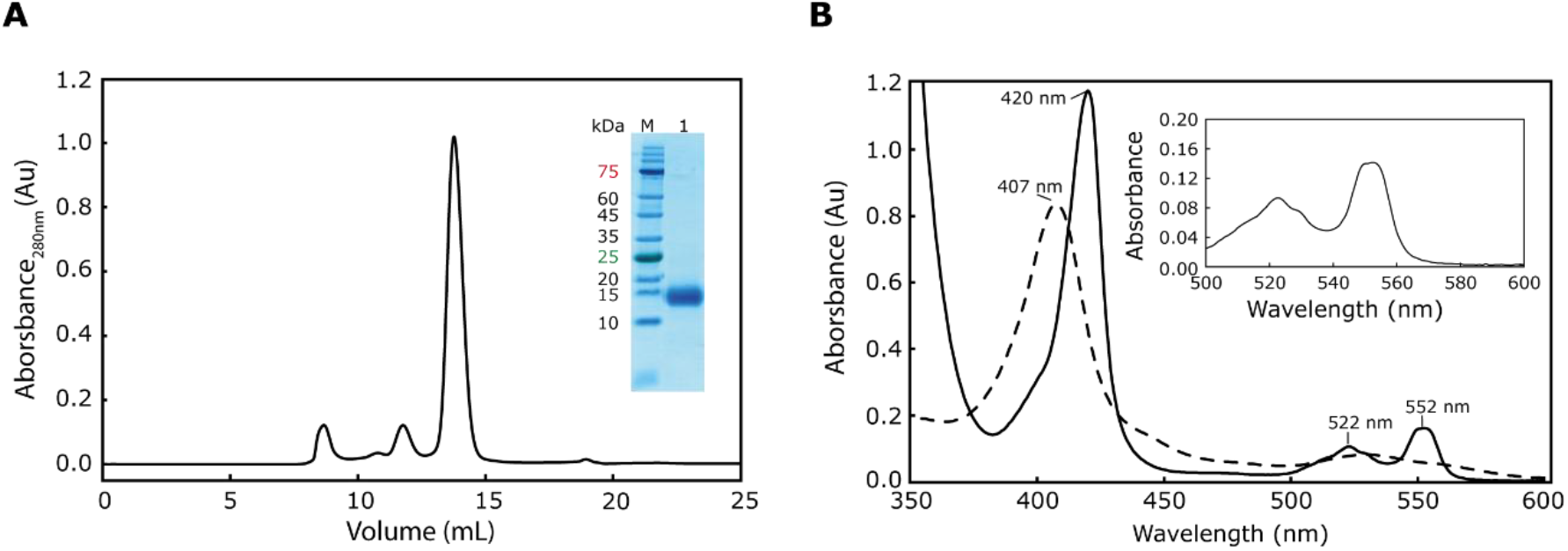
CbcS purification and UV-Visible spectroscopic characteristics. **(A)** Size-exclusion chromatography profile from the final step of CbcS purification. The inset displays an SDS-Page gel showing the purified CbcS protein (lane 1) together with the molecular weight marker (lane M). **(B)** UV-Visible spectra of CbcS (1.5 µM in 10 mM phosphate buffer, pH 8) recorded at room temperature. The dashed line corresponds to the oxidized state and the solid line represents the reduced form. The inset displays a zoom of the reduced spectrum between 500 and 600 nm showing a splitting of the characteristic peak of *c*-type cytochromes at 552 nm.

Furthermore, the N-terminal Strep-tag was cleaved, producing CbcS_cyt_, and 1D ^1^H NMR analysis also revealed that the presence of the Strep-tag does not affect the protein fold since the hemes’ resonances are unaltered (Figure S3). This result validates the usage of ST-CbcS_cyt_ as a representative model for CbcS_cyt_.

The presence of four heme groups in the produced proteins was confirmed by the pyridine hemochromogen assay and by mass spectrometry, and the molar extinction coefficient for CbcS was determined by the Lowry colorimetric assay at 552 nm in the reduced for (ε_552nm_ = 96.9 mM^-1^.cm^-1^).

### CbcS shows a split α-band in the reduced state

UV-visible spectra were acquired in both oxidized and reduced states and show typical characteristics of low-spin *c*-type hemes: a *Soret* band that shifts from 407 nm in the oxidized state to 420 nm when reduced and an α-band and β-band at 552 nm and 522 nm, respectively, in the reduced state (Figure 2B). Intriguingly, CbcS displays an interesting feature, since it shows a wider α-band with some splitting. This was only reported for mono and diheme cytochromes ^49^ and is probably associated with the relative arrangement between the hemes and its axial ligands, since the splitting disappeared in the pyridine-hemochromogen assay (data not shown).

CbcS heme groups’ spin state was evaluated in 1D ^1^H NMR spectra (Figure S4). These spectra provide valuable information on the spin state and axial ligands of heme groups ^50^. The signals for low- and high-spin hemes have some distinctive features in the reduced and oxidized state. In the oxidized state high-spin cytochromes exhibit broad heme methyl signals above 40 ppm, whereas low-spin cytochromes typically show resonances up to 40 ppm. In the reduced state, high-spin cytochromes display signals over a broad range (30 to -15 ppm), while low-spin cytochromes exhibit signals within a narrower range (10 to -5 ppm). The 1D ^1^H NMR spectra of CbcS in its oxidized and reduced states exhibit distinct differences, reflecting changes in the heme environment. In the reduced state, signals between 10 and -5 ppm confirm low-spin hemes, while the oxidized state displays broader peaks from 40 to -5 ppm due to paramagnetic effects. The absence of characteristic methionine signals supports a bis-histidine coordination for all four hemes ^51,52^. Some expected heme methyl signals are likely broadened beyond detection due to unpaired electrons, a phenomenon observed in other multiheme cytochromes.

### NMR validates the hemecore predicted by AlphaFold

CbcS has a predicted hemecore with the four hemes arranged in a chain-like architecture, with the meso protons 20H facing each other in pairs, heme I with heme II and heme III with heme IV (Figure 3A). When compared with the arrangement of STC hemes, CbcS hemes I/II/III superimpose with hemes II/III/IV of STC.

**Figure 3.**
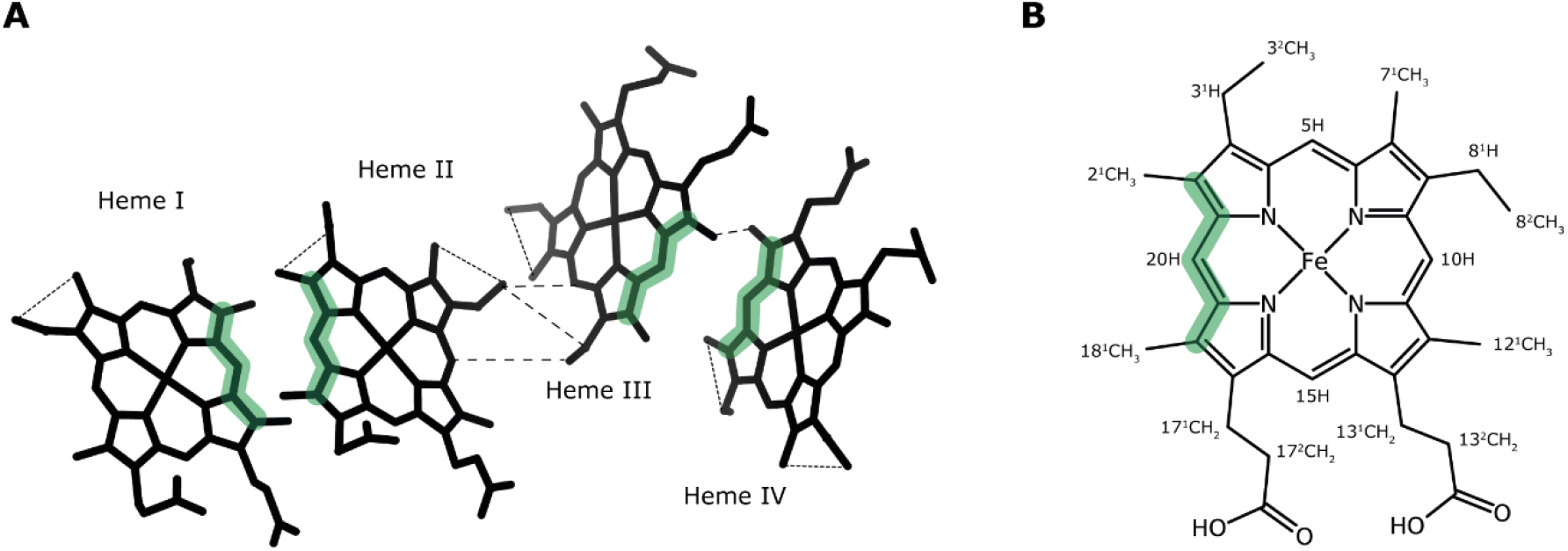
NOE connectivities observed for CbcS in the reduced state. **(A)** NOE connectivities observed in the 2D ^1^H NOESY spectra of CbcS represented in the hemecore predicted by AlphaFold. Large dashed and small dashed lines represent inter-heme connectivities and intra-heme connectivities between different heme faces, respectively. **(B)** Diagram of heme *c* with numbering according to the IUPAC-IUB nomenclature ^58,59^. In both panels, the heme faces of meso proton 20H are highlighted in green.

CbcS hemecore arrangement was analyzed by acquiring 2D NMR homonuclear experiments in the fully reduced state, specifically 2D ^1^H TOCSY and 2D ^1^H NOESY. Total Correlation Spectroscopy (TOCSY) utilizes scalar coupling to establish correlations between homonuclear spins within a spin system. Regarding heme resonances, each heme only gives rise to two cross-peaks corresponding to connectivities between 3^1^H/8^1^H (∼6 ppm) and 3^2^CH_3_/8^2^CH_3_ (∼1.5-3 ppm) (Figure 3B). The TOCSY spectra of CbcS shows eight signals, further confirming the correct assembly of four heme groups (Figure S5). These eight resonances are also the starting point to assign the resonances to specific heme groups.

The 2D ^1^H NOESY spectrum provides information on spatial distance between spins since Nuclear Overhauser Effect Spectroscopy (NOESY) relies on dipolar interactions to correlate spins within a proximity of up to 5 Å ^53^. In the fully reduced 2D ^1^H NOESY spectra of cytochromes, specific heme substituents display signals in distinct spectral: meso protons (5H, 10H, 15H, 20H) have chemical shifts between 8–10 ppm, thioether methines (3^1^H, 8^1^H) between 6–7 ppm, methyl groups (2^1^CH_3_, 7^1^CH_3_, 12^1^CH_3_, 18^1^CH_3_) between 3–4.5 ppm, and thioether methyls (3^2^CH_3_/8^2^CH_3_) between -1–2.5 ppm.

After the identification of the thioether connectivities in the TOCSY spectrum, the next step for heme resonance assignment was the identification of the meso protons 5H/10H that bind to it and the corresponding methyl groups, thus identifying each 5H/10H face of the heme, two per heme. Once these connectivities were established, the 20H, 2^1^CH_3_, and 18^1^CH_3_ signals were identified. Curiously, the meso protons 20H and heme methyls 2^1^CH_3_ and 18^1^CH_3_ exhibit upfield shifts (around 7 ppm) when compared to cytochrome *c*_3_ (around 8-10 ppm as the other meso protons), and as observed for STC hemes II and III, that also have the 20H facing each other ^41^. After identification of these heme faces, the next step was to connect the faces within each heme and then identify inter-heme connectivities. However, there were several superimpositions between the signals of the different heme substituents which made this task more difficult, even after acquiring spectra at different temperatures. In fact, the attribution of specific resonances to the corresponding heme groups started by identifying the two aromatic residues in the polypeptide chain (Tyr63 and Tyr159) (Figure S5). NOE cross peaks were observed between the signal at 7.05 from one tyrosine side chain and one 5H/10H and 3^2^CH_3_/8^2^CH_3_ face. As Tyr63 is close to heme I in the AlphaFold model, this specific face was identified as 5H/7^1^CH_3_/3^2^CH_3_/3^1^H signals of heme I. Subsequent analysis focused on identifying heme substituents with connectivities across different faces of the same heme or between adjacent hemes. The connectivities that led to the heme assignment, and that confirm the heme arrangement predicted by AlphaFold, are presented in Figure 3. The heme resonance assignment proposed for CbcS at different temperatures is presented in Table S2.

### Heme IV is CbcS’ electron exit gate

In tetraheme cytochromes four consecutive reversible steps of one-electron convert the fully reduced to the fully oxidized state, defining five different redox stages each containing microstates with the same number of oxidized hemes ^54^. As the chemical shifts of the heme signals are different in the reduced and oxidized states (Figure S4), when fast intramolecular electron exchange (between different microstates within the same oxidation stage) and slow intermolecular electron exchange (between different oxidation stages) relative to the NMR timescale is achieved, the individual heme substituent signals can be discriminated in the different oxidation stages. The acquisition of 2D EXSY NMR experiments allow thus to monitor the individual oxidation patterns of the hemes.

2D EXSY NMR redox titrations were acquired for CbcS at 1mM and initially at 25°C to mimic the ideal conditions observed for other tetraheme cytochromes, as STC ^41^. However, broad signals, in some oxidation stages even beyond detection, were obtained. Consequently, temperature was lowered to 16°C, and then further to 9°C to decrease signals broadness (Figure S6). Several 2D EXSY spectra were then obtained at different oxidation stages at 9°C, corresponding to each 1D ^1^H spectrum presented in Figure S6, and analyzed. Spectra in early oxidation stages allowed to identify cross peaks for stage 0-1 for all four methyls of heme IV (Figure S8), and for some, it was possible to monitor all five stages of oxidation (Table S3). However, the same monitorization was not possible for the other three hemes, since only cross peaks for intermediate stages were observed, not allowing to correlate with the assignment in the full reduced state. Usually, it is also possible to attribute redox patterns to specific hemes after assigning the heme resonances in the fully oxidized state ^55^. Nevertheless, different conditions (as sample concentration and temperature) were tested for acquisition of 2D spectra (NOESY, TOCY, HMQC) but no signals were obtained for resonances downfield of the spectra above ∼28ppm (data not shown). Further conditions, such as acquisition at different pH values and different magnetic fields should be evaluated. Anyhow, it was possible to determine that heme IV is the first to oxidize, and thus, this heme is the exit gate for electrons in Cbc4.

### The redox active windows of CbcS and its putative redox partners overlap

The UV-visible spectra of heme proteins also exhibit distinct spectroscopic properties, including characteristic bands in both their reduced and oxidized states (Figure 2B), which enables monitoring their redox state during potentiometric titrations. In this study, redox titrations monitored by visible spectroscopy were performed on ST-CbcS_cyt_ at two different pH values to determine its apparent reduction potential (*E*_*app*_) and working potential range. The *E*_*app*_ values were determined to be -139 ± 3 mV at pH 7 and -151 ± 1 mV at pH 8 (Figure 4 and Table S4). The observed pH dependence of the reduction potential can be attributed to electrostatic effects, as the deprotonation of nearby acidic or basic groups reduces the heme’s affinity for electrons, leading to a decrease in the reduction potential. This difference in reduction potential across pH values highlights the redox-Bohr effect in CbcS. To understand the redox characteristics of CbcS in the context of *G. sulfurreducens* EET networks, its redox active window was compared with those of periplasmic triheme cytochromes from the PpcA family, which are potential redox partners of CbcS, and known terminal reductases (Figure 4 inset and Table S5). The reduction potential values obtained for CbcS were found to be only slightly more negative at both pH values compared to those reported for the PpcA family (except for PpcC at pH 7). This indicates that electron transfer to most cytochromes in the PpcA family is thermodynamically favorable, although it is less favorable when compared to other inner membrane oxidoreductases, such as CbcL and ImcH ^56,57^. The redox-active window of CbcS overlaps with that of the PpcA-E family, which may impact electron transfer. Therefore, the electron transfer reaction between CbcS and PpcA was monitored using NMR.

**Figure 4.**
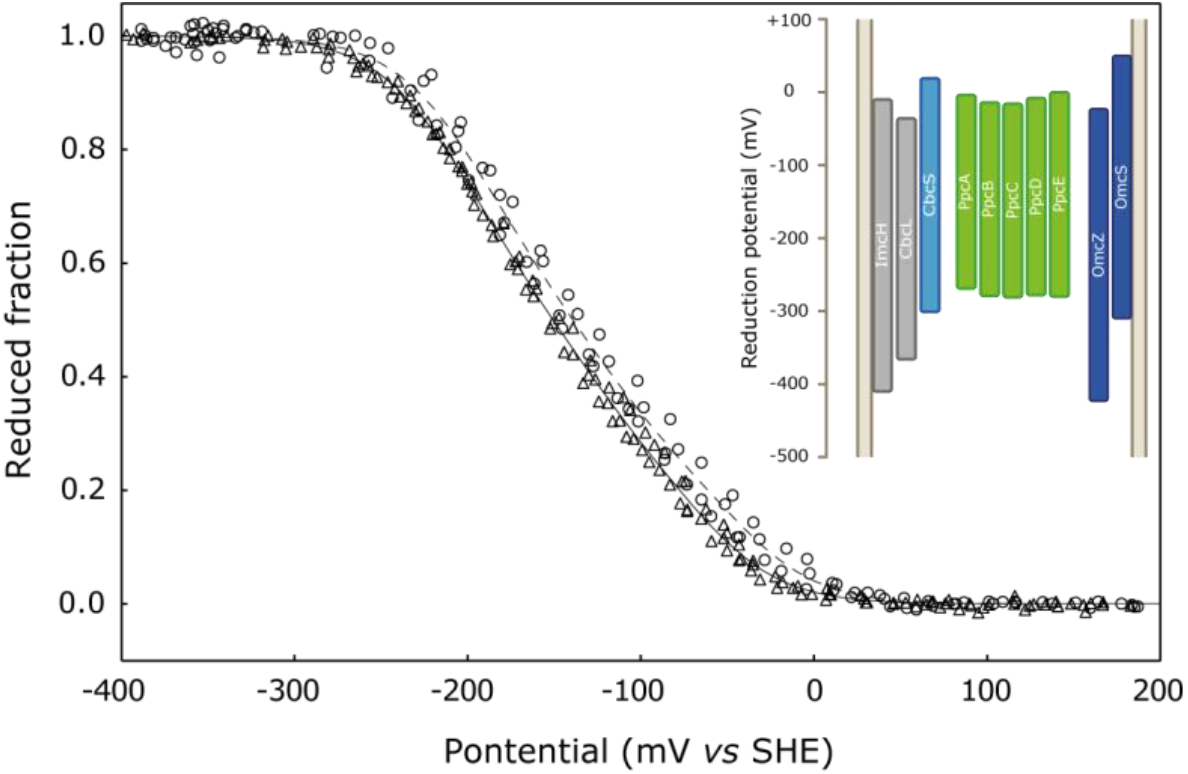
Redox characterization of CbcS. Potentiometric redox titrations of CbcS acquired with 8 µM of CbcS in 80 mM sodium phosphate buffer with NaCl (I = 250 mM) at 16°C. The experimental points at pH 8 and pH 7 are represented as triangles and circles, respectively. The solid line represents the fitted curve for pH 8 and the dashed line represents the fitted curve for pH 7. The inset shows the redox-active windows (1-99%) of *G. sulfurreducens* inner membrane quinone oxidases, periplasmic cytochromes and outer membrane reductases at pH 8 ^40,56,57,60–62^.

### CbcS in the EET framework of *G. sulfurreducens*

In the Cbc4 inner membrane quinone oxidase complex, the iron-sulfur protein CbcT likely transfers electrons from the menaquinone pool to CbcS, which then passes them to a redox partner in the periplasm. To investigate putative redox partners in the periplasm, electron transfer between reduced CbcS and oxidized PpcA was examined using NMR (Figure 5). The NMR experiment started with CbcS in its reduced state followed by titration with increasing heme equimolar amounts of oxidized PpcA. The additions of PpcA were done inside an anaerobic glovebox so that any changes were due to electron transfer between the proteins and not to the presence of oxygen ^42^. Following the first addition of oxidized PpcA, broad signals appeared at ∼17 ppm and ∼27 ppm. An initial observation let to conclude that the latter signal had to be attributed to CbcS, since PpcA does not show any resonances above 22 ppm. Later, these signals were respectively attributed to heme IV methyls 7^1^CH_3_ and 18^1^CH_3_ at oxidation stage 1 (Figure S8). This further confirms that heme IV is the exit gate for electrons even when electrons a transferred to a redox partner. As more oxidized PpcA was added, additional low-field resonances were observed, indicating that neither cytochrome was fully reduced nor oxidized. To verify that the two cytochromes were in an intermediate oxidation state, the NMR tube was opened and exposed to atmospheric O_2_, leading to the fully oxidation of CbcS and PpcA. The results obtained were not unexpected due to the total overlap between the redox active windows of CbcS and PpcA and was also previously observed for the interaction between CbcL or ImcH with PpcA ^56,57^. This superimposition justifies the fact electrons are not completely transferred from CbcS to PpcA, which can enable a continuous flow of electrons whenever the Cbc4 complex is reduced by the quinone pool and PpcA is oxidized by its upstream redox partner.

**Figure 5.**
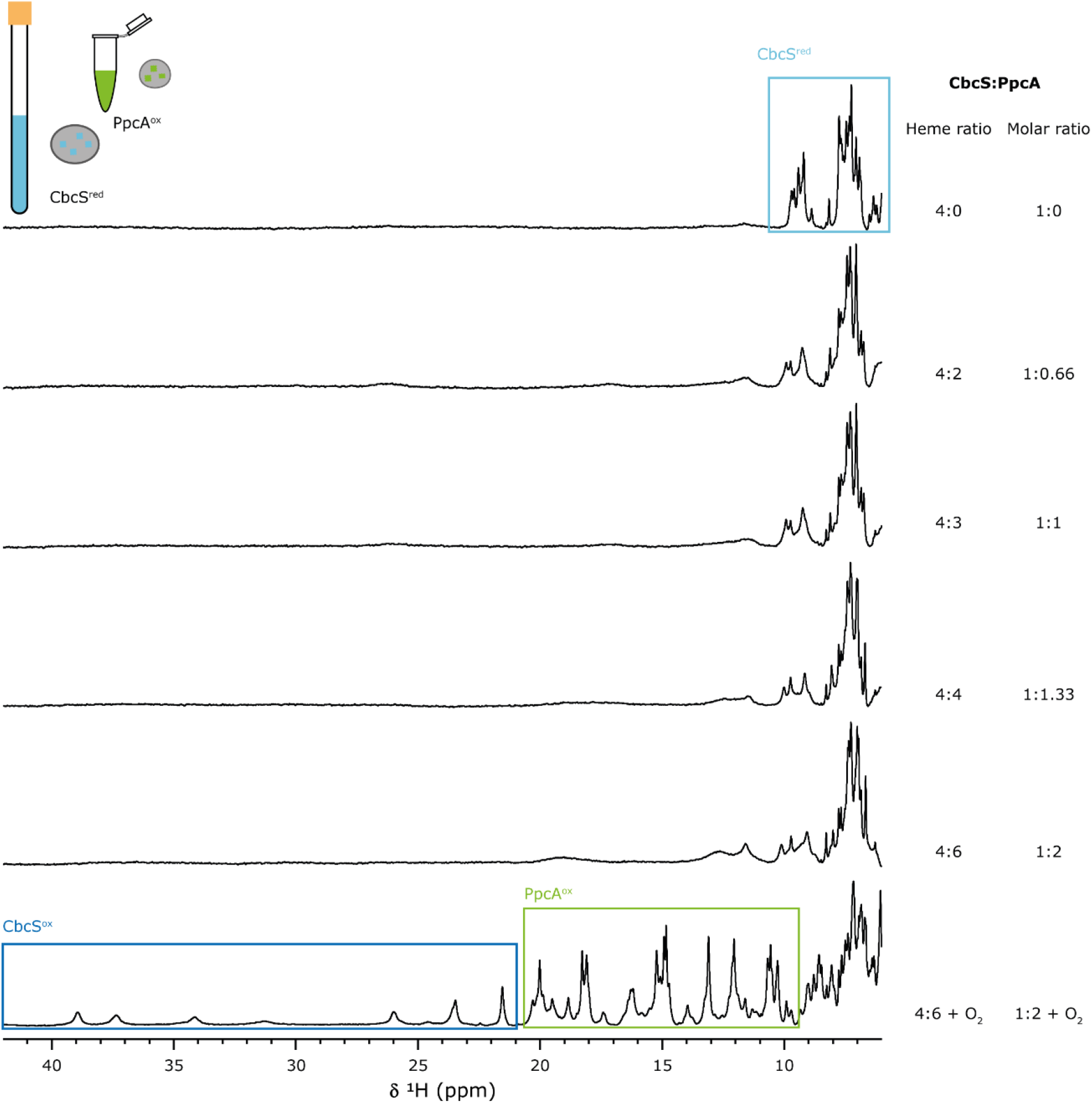
Electron transfer reaction between CbcS and PpcA monitored by NMR. Spectra were acquired at 20°C with 45 µM CbcS in 10 mM sodium phosphate pH 8 buffer with increasing equimolar amounts of PpcA, up to a 1:2 ratio (4:6 hemes ratio). The rectangles highlight the distinct spectral regions representing the cytochromes in their reduced and oxidized states.

## Conclusions

The knowledge on the metabolism of electroactive bacteria has increased in the past decades. Besides the importance of enhancing the capability of *Geobacter* for biotechnological applications, understanding the detailed molecular mechanisms for EET is essential. This works establishes groundwork on the characterization of CbcS, a multiheme cytochrome that is part of an inner membrane respiratory complex that is thought to act as a quinone-cytochrome *c* oxidoreductase ^13^. The gene coding for CbcS was upregulated when *G. sulfurreducens* was grown on Fe(III) oxides but when *cbcS* gene was deleted, mutant strains grew at wild-type levels, suggesting that CbcS is not essential for Fe(III) or Mn(IV) reduction ^12^. However, only CbcS was mutated and not the complete Cbc4 complex, so the FeS cluster protein CbcT could be transferring electrons to other cytochromes in the periplasm. Further research is essential to check the role of Cbc4 complex and its involvement in parallel pathways for EET. This work provides initial biochemical data on the properties of CbcS and provides relevant information for a new protein of the NapC/NirT family, that was not possible to obtain for example for the homologous cytochrome CymA from *Shewanella* species. The determined redox properties, and interaction studies with its putative redox partner in the periplasm, presents a possible new route for electron transfer, and places CbcS within the broader EET framework of *G. sulfurreducens*.

## Acknowledgments

This work was supported by Fundação para a Ciência e a Tecnologia (FCT, Portugal) through grants 2022.11900.BD (JMAA), PTDC/BIA-BQM/4967/2020 (CAS), EXPL/BIA-BQM/0770/2021 (LM), UIDP/04378/2020 and UIDB/04378/2020 (UCIBIO – Research Unit on Applied Molecular Biosciences), and LA/P/0140/2020 (i4HB – Associate Laboratory Institute for Health and Bioeconomy). The NMR spectrometers are part of the National NMR Network and are supported by FCT (ROTEIRO/0031/2013 and PINFRA/22161/2016) cofounded by FEDER through COMPETE 2020, POCI, PORL and FCT through PIDDAC.

