## Supplementary Information for "Preliminary characterization of CbcS from *Geobacter sulfurreducens*’ Cbc4 complex: a putative novel respiratory pathway"

### Supplementary Tables

**Table S1. Sequence of the primers used.**

| Primer | Sequence |
|---|---|
| **ST-CbcS_FW30** | GGAGTCATCCTCAATTCGAAAAATCCGGAGATTTCGGGGTGCATGAGC |
| **ST-CbcS_RV** | GCCATTTTTCACTTCACAGGTCAAGCTTTTATTCATAGCGCTTCAACTCCTC |
| **ST-CbcS_FW59** | GGAGTCATCCTCAATTCGAAAAATCCGGACGGACCAAAGCCTACTGC |
| **ST-CbcS59-TEVfw** | ATTTTCAGTCTCGGACCAAAGCCTAC |
| **pVA203-ST-insertTEV_rv** | ACAGGTTTTCTCCGGATTTTTCGAATTG |

**Table S2. Chemical shifts (ppm) of the heme substituents of CbcS in the reduced state at pH 8 and 9°C ,16°C and 25°C.**

| | Temperature (°C) | Heme I | Heme II | Heme III | Heme IV |
|---|---|---|---|---|---|
| **5H** | 9 | 9.21 | 9.22 | 9.77 | 9.41 |
| | 16 | 9.20 | 9.23 | 9.77 | 9.41 |
| | 25 | 9.20 | 9.21 | 9.76 | 9.42 |
| **10H** | 9 | 9.30 | 8.88 | 9.42 | 9.20 |
| | 16 | 9.30 | 8.87 | 9.42 | 9.20 |
| | 25 | 9.26 | 8.87 | 9.40 | 9.19 |
| **20H** | 9 | 7.34 | 7.65 | 7.72 | 7.55 |
| | 16 | 7.34 | 7.64 | 7.74 | 7.56 |
| | 25 | 7.33 | 7.65 | 7.72 | 7.55 |
| **$2^1CH_3$** | 9 | 0.94/3.46 | 1.01 | 2.88 | 2.63 |
| | 16 | 0.92/3.40 | 0.99 | 2.87 | 2.63 |
| | 25 | 0.92/3.46 | 0.96 | 2.88 | 2.74 |
| **$7^1CH_3$** | 9 | 2.79 | 3.04 | 4.00 | 3.65 |
| | 16 | 2.82 | 3.05 | 4.00 | 3.65 |
| | 25 | 2.83 | 3.04 | 3.99 | 3.65 |
| **$12^1CH_3$** | 9 | 3.48 | 3.48 | 3.58 | 3.51 |
| | 16 | 3.48 | 3.49 | 3.59 | 3.51 |
| | 25 | 3.48 | 3.379 | 3.57 | 3.51 |
| **$18^1CH_3$** | 9 | 0.94/3.46 | 2.72 | -0.80 | 1.24 |
| | 16 | 0.92/3.40 | 2.75 | -0.81 | 1.25 |
| | 25 | 0.92/3.46 | 2.74 | -0.82 | 1.21 |
| **$3^1H$** | 9 | 6.05 | 6.21 | 6.41 | 6.35 |
| | 16 | 6.06 | 6.21 | 6.41 | 6.34 |
| | 25 | 6.06 | 6.21 | 6.41 | 6.34 |
| **$8^1H$** | 9 | 6.33 | 6.01 | 6.25 | 6.00 |
| | 16 | 6.32 | 5.99 | 6.25 | 6.00 |
| | 25 | 6.30 | 5.99 | 6.23 | 6.00 |
| **$8^2CH_3$** | 9 | 2.24 | 1.50 | 2.51 | 2.16 |
| | 16 | 2.22 | 1.50 | 2.50 | 2.16 |
| | 25 | 2.23 | 1.50 | 2.51 | 2.14 |
| **$3^2CH_3$** | 9 | 2.18 | 1.93 | 2.77 | 2.08 |
| | 16 | 2.17 | 1.97 | 2.76 | 2.08 |
| | 25 | 2.16 | 1.94 | 2.74 | 2.08 |

**Table S3. Redox dependence of CbcS heme IV methyl chemical shifts and heme oxidation fractions at pH 8.1 and 9°C).** *nd* – not determined.

| Oxidation stage | Chemical shift (ppm) | | | | Oxidation fraction | | | |
|---|---|---|---|---|---|---|---|---|
| | $2^1CH_3^{IV}$ | $7^1CH_3^{IV}$ | $12^1CH_3^{IV}$ | $18^1CH_3^{IV}$ | $2^1CH_3^{IV}$ | $7^1CH_3^{IV}$ | $12^1CH_3^{IV}$ | $18^1CH_3^{IV}$ |
| **0** | 2.63 | 3.65 | 3.51 | 1.24 | - | 0 | - | 0 |
| **1** | 7.62 | 17.06 | 11.60 | 26.62 | - | 0.65 | - | 0.65 |
| **2** | - | 19.54 | - | 31.10 | - | 0.77 | - | 0.76 |
| **3** | - | 23.72 | - | 39.43 | - | 0.98 | - | 0.97 |
| **4** | - | 24.18 | - | 40.57 | - | 1 | - | 1 |

**Table S4. CbcS macroscopic reduction potentials and apparent midpoint potentials (*vs* SHE) at pH 7 and pH 8 and 15°C.**

| | $E_1$ (mV) | $E_2$ (mV) | $E_3$ (mV) | $E_4$ (mV) | $E_{app}$ (mV) |
|---|---|---|---|---|---|
| **pH 7** | -212 ± 3 | -168 ± 3 | -112 ± 3 | -41 ± 3 | -139 ± 3 |
| **pH 8** | -222 ± 1 | -179 ± 1 | -123 ± 1 | -60 ± 1 | -151 ± 1 |

**Table S5. Apparent midpoint reduction potential values for CbcS, PpcA-family cytochromes (PpcA, PpcB, PpcC, PpcD and PpcE) [40,60], CbcL [56] and ImcH [57] at pH 7 and 8 and 15°C.**

| Cytochrome | $E_{app}$ pH 7 (mV) | $E_{app}$ pH 8 (mV) |
|---|---|---|
| **CbcS** | -139 | -151 |
| **CbcL** | - | -194 |
| **ImcH** | - | -196 |
| **PpcA** | -117 | -138 |
| **PpcB** | -137 | -146 |
| **PpcC** | -143 | -146 |
| **PpcD** | -132 | -148 |
| **PpcE** | -134 | -138 |

### Supplementary Figures

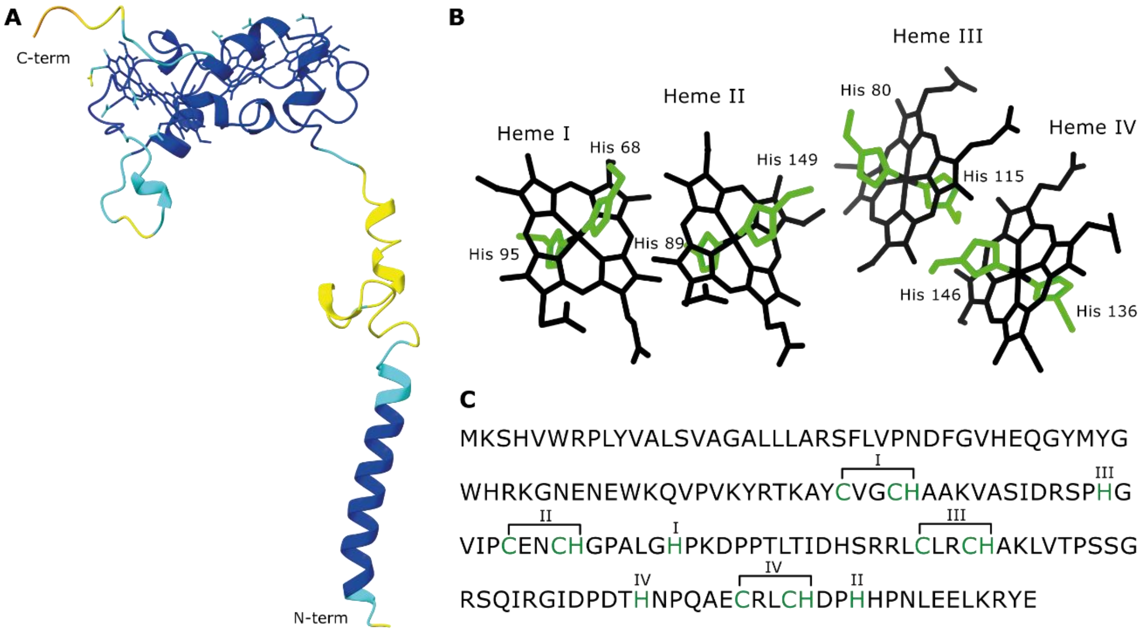


**Figure S1. CbcS structural model and sequence analysis. (A)** CbcS AlphaFold model showed with the pLDDT confidence of the model. The residues in blue have very high confidence (pLDDT > 90), in cyan have high confidence (90 >pLDDT > 70), in yellow low confidence (70 >pLDDT > 50) and in orange very low confidence (pLDDT < 50). **(B)** CbcS hemecore arrangement predicted by AlphaFold with heme axial ligands in green. **(C)** CbcS protein sequence. The letters in green highlight the CXXCH binding motifs and the axial histidine ligands. Each binding motif and axial histidine is identified to which heme it corresponds to.

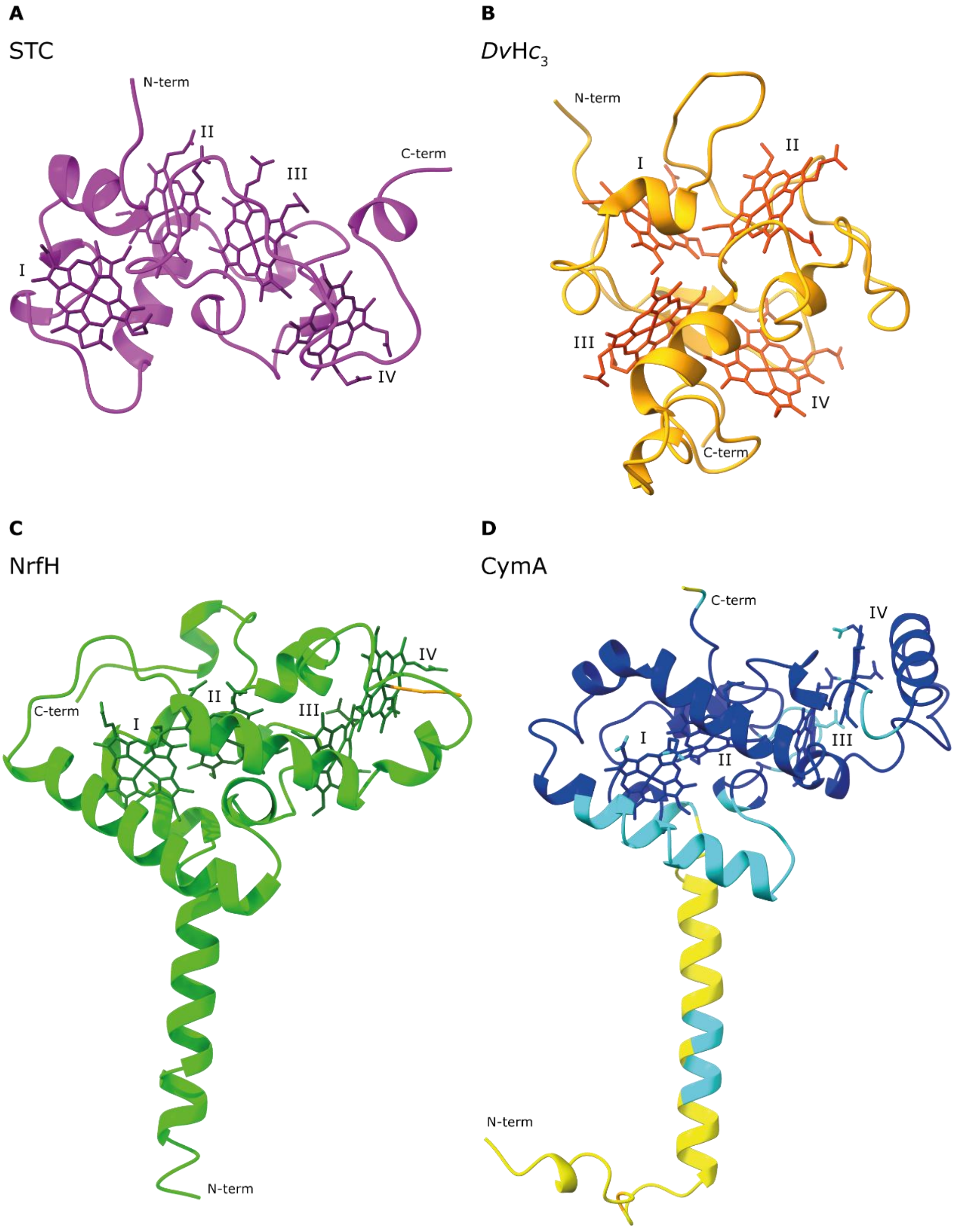


**Figure S2. Structure of different tetraheme proteins. (A)** Solution NMR structure of small tetraheme cytochrome c (STC) from *Shewanella frigidimarina* (PBD 2K3V) [43]. **(B)** Solution NMR structure of cytochrome $c_3$ from *Desulfovibrio vulgaris* (Hildenborough) (PBD 1A2I) [44]. **(C)** Crystal structure of cytochrome *c* nitrite reductase NrfHA complex from *Desulfovibrio vulgaris* (PBD 2J7A) [45]. The residue displayed in orange is the NrfH heme IV distal ligand, a lysine from NrfA. **(D)** AlphaFold predicted model for CymA from *S. frigidimarina*, colored by pLDDT confidence. The residues in blue have very high confidence (pLDDT > 90), in cyan have high confidence (90 >pLDDT > 70), in yellow low confidence (70 >pLDDT > 50) and in orange very low confidence (pLDDT < 50).

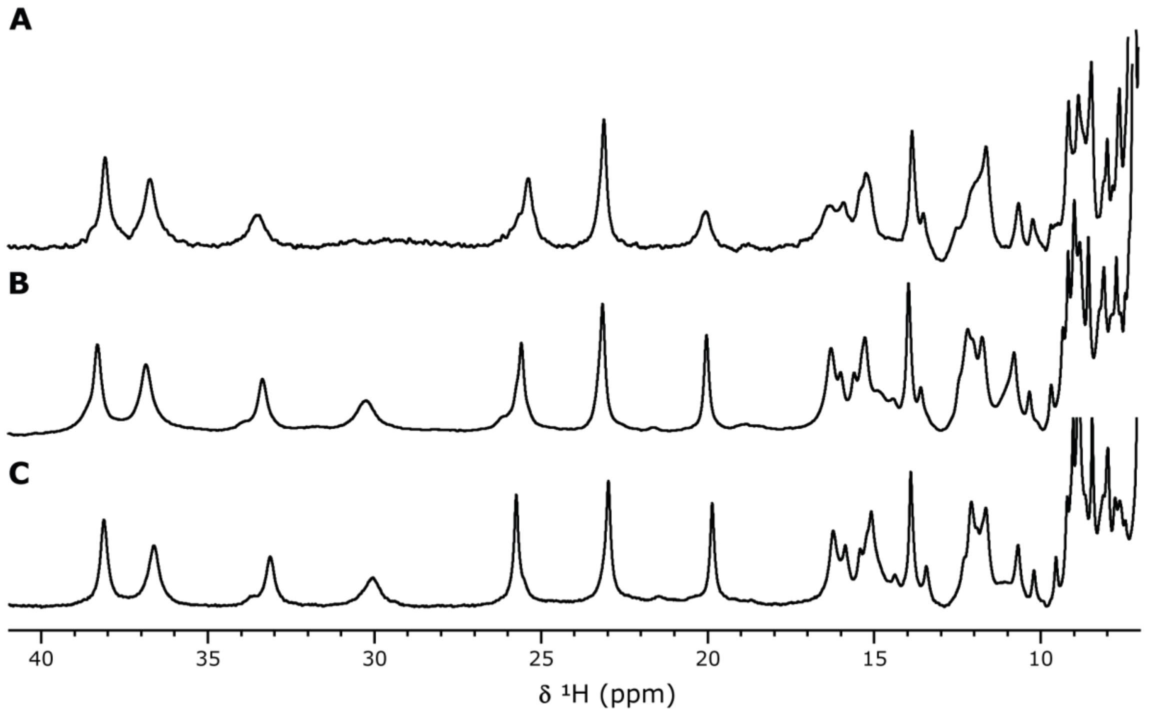


**Figure S3. 1D $^1$H NMR spectra of CbcS constructs in the oxidized state. (A)** ST-CbcS$_{pp}$ ~100 µM in 50 mM sodium phosphate pH 8. **(B)** ST-CbcS$_{cyt}$ 300 µM in 80 mM sodium phosphate pH 8 (I = 250mM). **(C)** CbcS$_{cyt}$ 1 mM in 80 mM sodium phosphate pH 8 (I = 250mM). All spectra were acquired at 25°C in $^2H_2O$.

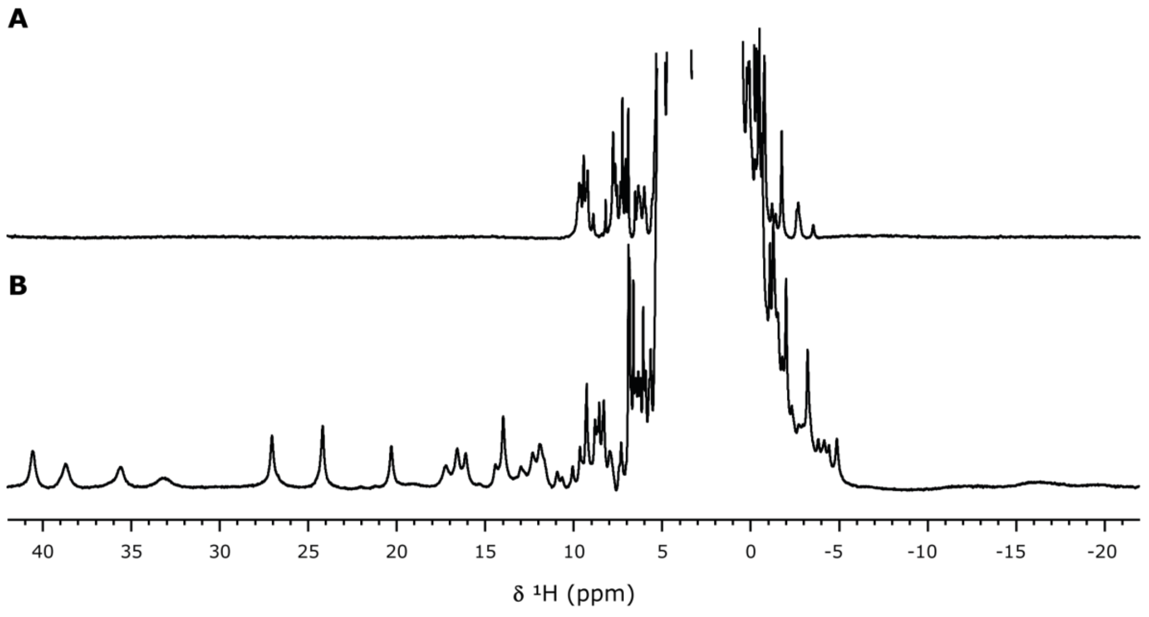


**Figure S4. 1D $^1$H NMR spectra of CbcS in the reduced (A) and oxidized (B) states.** Spectra were acquired at 9°C with 1mM CbcS in 80 mM sodium phosphate with NaCl (I= 250 mM) pH 8.1 in $^2H_2O$.

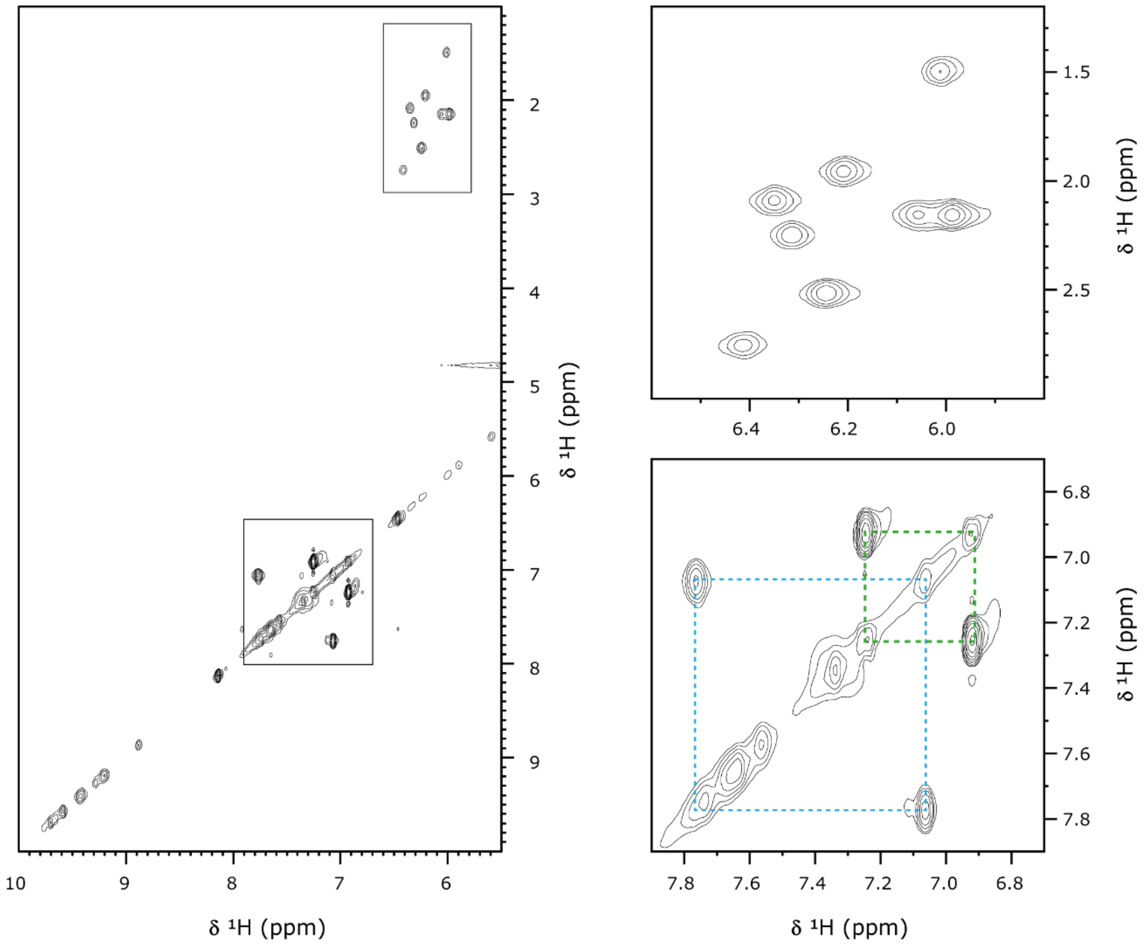


**Figure S5. TOCSY connectivities of CbcS.** Low field region of the 2D $^1H$ TOCSY spectrum of CbcS highlighting the heme thioether signal and aromatic side chains (left side). On the right, the top inset is a zoom of the connectivities between thioether protons $3^1H/8^1H$ and thioether methyls $3^2CH_3/8^2CH_3$ and the bottom inset is a zoom of the signals from the side chain of aromatic residues Tyr63 (blue) and Tyr159 (green). The spectrum was acquired at 25°C with 1 mM CbcS in 80 mM sodium phosphate pH 8.1 with NaCl (I = 250mM) in $^2H_2O$.

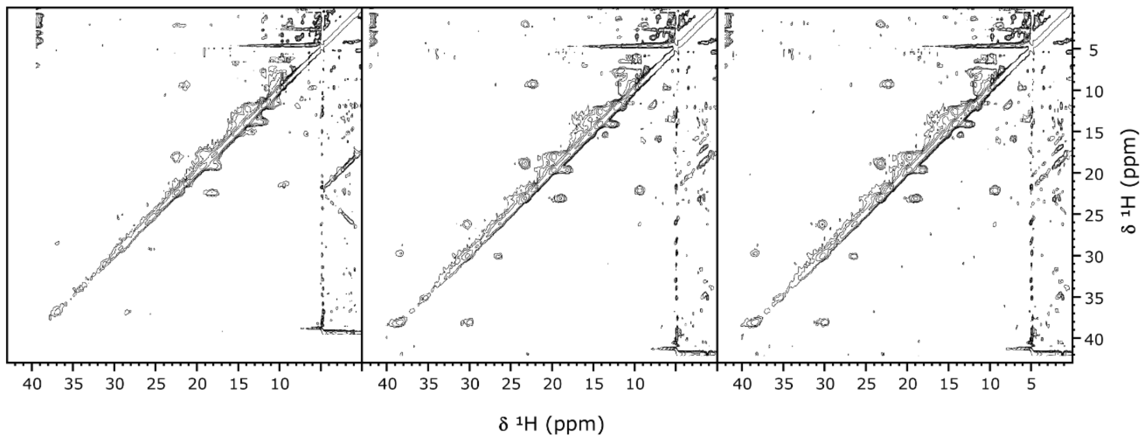


**Figure S6. 2D EXSY NMR spectra at different temperatures.** The spectra were acquired with 1mM CbcS in 80 mM sodium phosphate pH 8.1 with NaCl (I = 250mM) in $^2H_2O$ in the same oxidation state at 25°C (left), 16°C (middle) or 9°C (right)

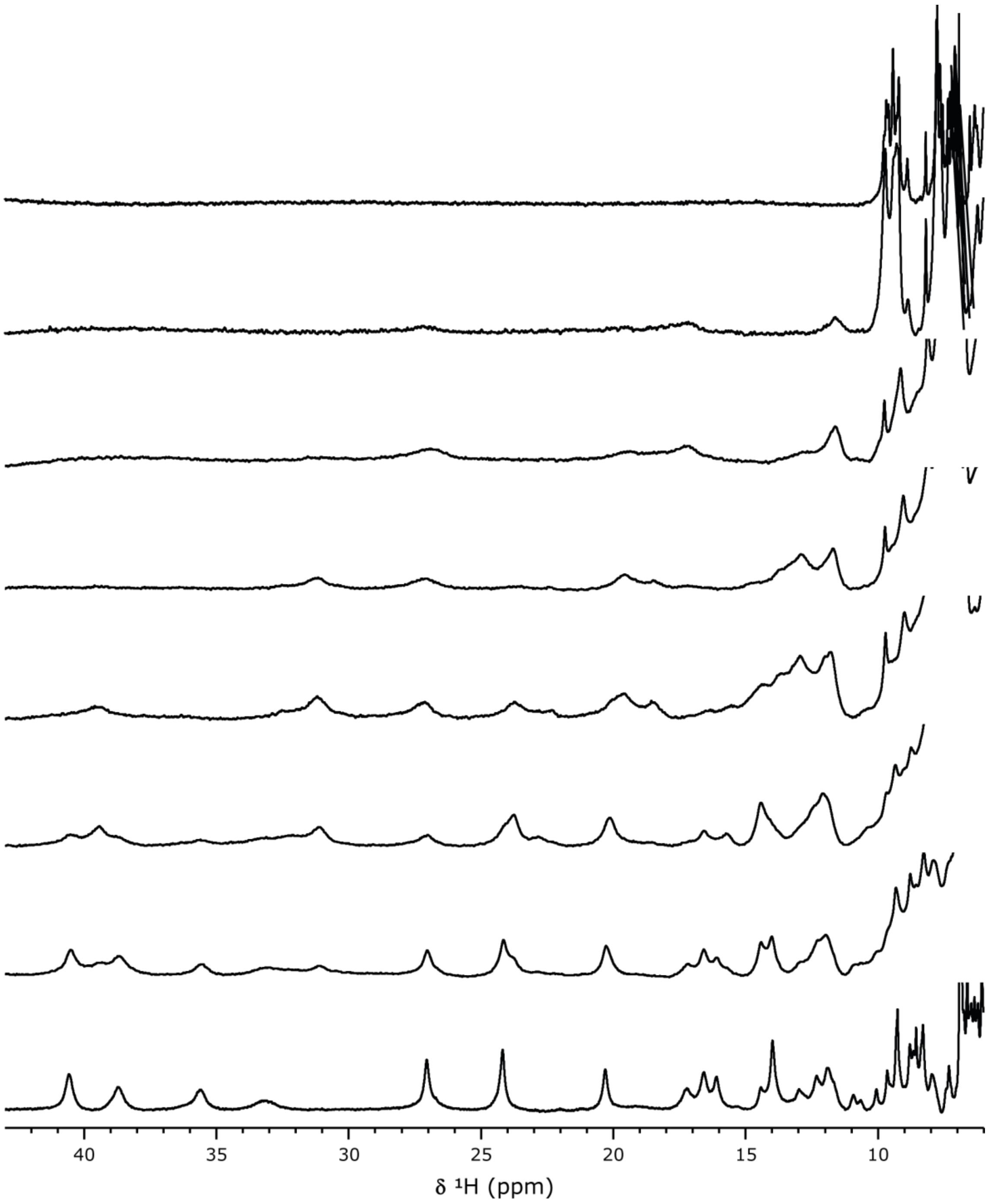


**Figure S7. 1D ¹H NMR spectra of CbcS at different redox states.** Spectra were acquired after addition of increasing amounts of $O_2$ at 25°C with 1mM CbcS in 80 mM sodium phosphate pH 8.1 with NaCl (I = 250mM) in $^2H_2O$.

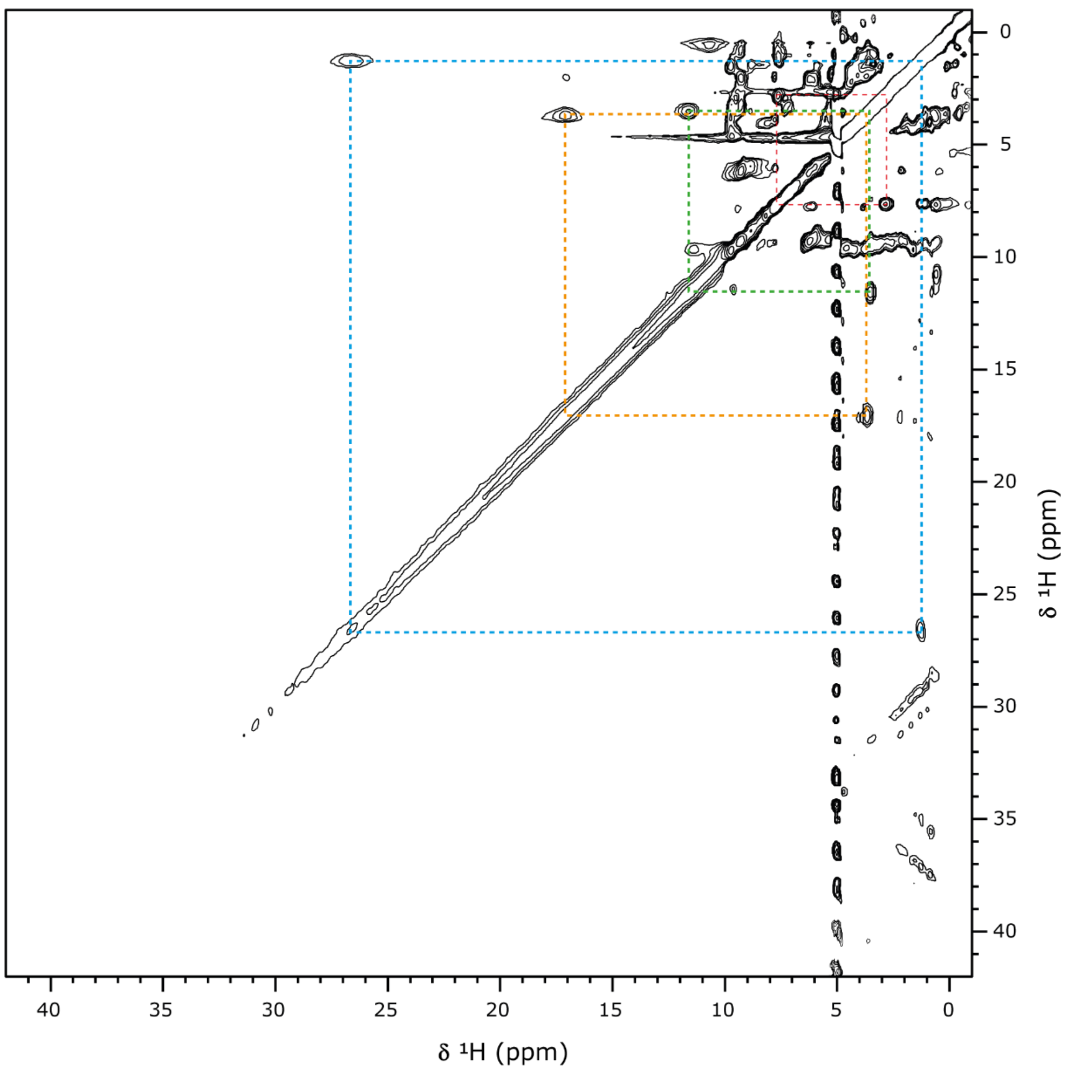


**Figure S8. 2D $^1$H EXSY NMR spectra of CbcS in an intermediate state.** The spectrum was acquired at 9°C with 1 mM CbcS in 80 mM sodium phosphate pH 8.1 with NaCl (I = 250mM) in $^2H_2O$ in an early oxidation stage (corresponding to the second 1D $^1$H spectrum from the top on Figure S7). The dashed lines connect each of the cross-peaks for the four methyl groups of heme IV in stage 0-1 ($2^1CH_3^{IV}$ in red, $7^1CH_3^{IV}$ in orange, $12^1CH_3^{IV}$ in green and $18^1CH_3^{IV}$ in blue), showing it is the first heme to oxidize.
